# Correlating stress reduction and eye movement patterns in a world famous Kyoto Japanese garden

**DOI:** 10.1101/2023.08.21.554170

**Authors:** Seiko Goto, Hiroki Takase, Keita Yamaguchi, Tomoki Kato, Minkai Sun, Aoi Koga, Liang Tiankai, Isamu A. Poy, Karl Herrup

## Abstract

Visual stimuli have been repeatedly shown to elicit significant non-visual responses. In a continuing effort to explore the unique effects of viewing a Japanese garden on the physiological and psychological metrics of stress, we gained access to the world famous Murin-an garden in Kyoto, Japan. This well-maintained observation garden was designed to be viewed while seated at a single vantage point to maximize the impact of the visual scene. As a control, we used a public garden on the campus of Kyoto University that was designed in a similar style. Sixteen college age students were asked to view both gardens while we monitored their pulse rates and tracked their eye movements. We used the POMS questionnaire to determine the effect of the garden viewing on the mood of the participants. We found that the Murin-an garden was more effective in decreasing pulse rate and improving mood than the University garden. The eye tracking data showed that during their Murin-an viewing the participants gaze ranged far more broadly across the visual field both the X-Y plane and in depth, and the speed with which the eyes moved from point-to-point was greater. Taken together, our data suggest that no one element in the garden was dominant in eliciting the changes in heart rate and mood. Rather, it was the breadth and rapidity of the shifts in gaze that drove the effects, a conclusion with implications for other interventions aimed stress reduction.

**Significance:** Views of nature and natural phenomena have a well-recognized calming effect on humans that has recognized therapeutic value in both medical and psychological settings. Our work explores the source of this effect by having participants view Murin-an, a world-famous Japanese style garden. Using both psychological and physiological measures, we confirm and extend earlier findings showing that a well constructed garden can effectively lower heart rate and improve mood within minutes. We also find, by analogy with eye movement desensitization and reprograming (EMDR), that it is participants’ rapidly shifting gaze rather than a single specific visual object is the most likely source of the calming effect.

## Introduction

A traditional style of a Japanese garden is known as an “observation garden.” Unlike most gardens where a visitor is expected to move through its space and appreciate its elements from many different visual perspectives, an observation garden is designed to be viewed while seated at a single vantage point. Indeed, the location of this vantage point is a critical design feature that is carefully chosen by the designer to maximize the impact of the visual scene. One reason for imposing this level of control on the viewer is that observation gardens were meant to be more than merely esthetically pleasing. For example, they were favored within Zen temples as aids for meditation and thus performed an important function for the temple residents. Today, in addition to their original functions, many observation gardens have become tourist attractions. While their esthetic beauty is part of their appeal, it is likely that another important draw is their documented history of providing a calming effect on the viewer. The centuries of landscape architects who perfected the design of the observation gardens no doubt honed their skills by trial and error. Yet from a modern perspective the consistent impact of their creations on a visitor raises questions as to what might be the underlying physiological and neurobiological mechanisms that these architects tapped into.

The effect of a natural environment on human physiology has been noted both anecdotally and in formal research studies. Ulrich was one of the first modern practitioners of the latter approach (Ulrich, 1984). His idea was that “most natural views apparently elicit positive feelings, reduce fear in stressed participants, hold interest, and may block or reduce stressful thoughts, they might also foster restoration from anxiety or stress.” Soon after this work was published, Kaplan & Kaplan in Attention Restoration Theory (ART), proposed that unconscious mild aesthetic experiences in nature can have a psychological healing effect (Kaplan, 1994; Kaplan and Kaplan, 1989). Other sensory stimulations have also been found to have positive health effects (Abraha et al., 2017; Blackburn and Bradshaw, 2014; Goris et al., 2016; Matthews, 2015; Moyle et al., 2014; Ohman et al., 2014; Vink et al., 2013). Works such as these open a critical window into the impact of the sensory environment on a person’s physiological state, yet most lacked a strong neurobiological explanation. As the field has progressed and imaging technology has become more widely used we now appreciate that receiving stimulation from a visual field such as a garden is routinely associated with activation of the specific regions of the brain.

Though the data is less extensive, there are increasing numbers of studies reporting on eye-movement activity during a visual stimulation involving a natural scene such as a garden. Our own work has focused on studies such as these. What we have repeatedly observed is that when an individual views a structured natural space, created using the design principles of a Japanese garden, they tend to take in the scene presented with more visual fixation points and longer fixation times, than when viewing other types of green spaces (Sun and Fujii, 2018). The parasympathetic nervous system, which is associated with relaxation, was also found to become dominant when the participants’ eyes move around a Japanese garden (Liu et al., 2020).

In the current study we sought to extend our work to the Murin-an garden in Kyoto Japan. Murin-an is one of the most famous Japanese gardens and has been designated as a historic site and place of scenic beauty by the Japanese government (Figure 1A). The garden, located in the heart of Kyoto, was originally built in 1894 as a villa for Yamagata Aritomo, the former prime minister of Japan. It was designed as an observation garden to be viewed from the center of the main room of the villa. The view takes advantage of a technique known as “borrowing scenery” by incorporating the nearby Mt. Higashi into the design. The garden is meant to represent a river flowing from a distant mountain, with a shallow pond and zig-zag stream surrounded by lush vegetation. This peaceful atmosphere has earned the garden a reputation as being one of the most relaxing in the region. We wished to ask how a viewer’s response to a classic observation garden such as this would compare with a Japanese style garden, but one that was not specifically designed as an observation garden.

**Figure 1.**
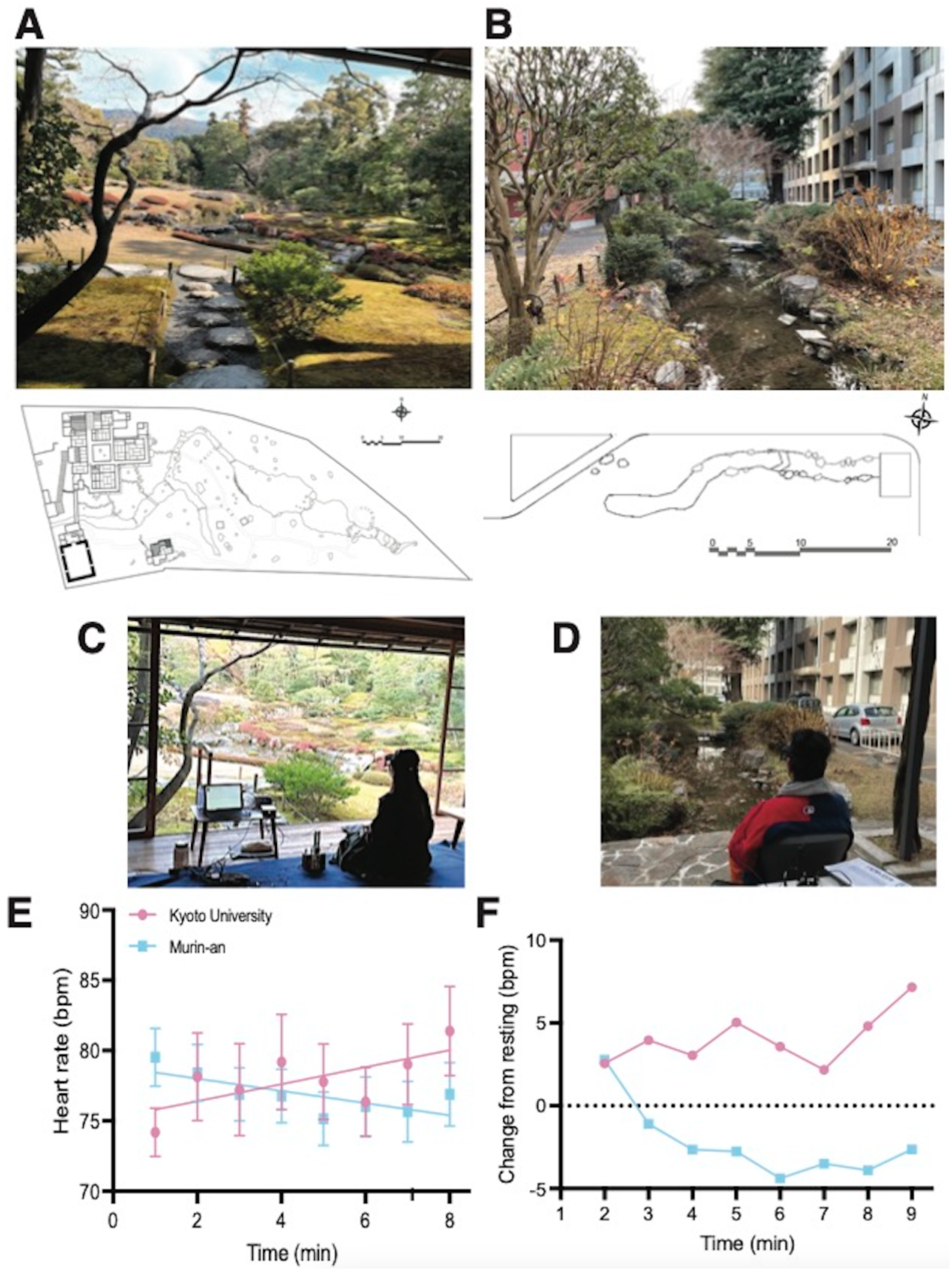
Photograph of the Murin-an garden on the day of the experiment. The drawing beneath the photo is the site plan as provided by Tomoki Kato. **B**. Photgraph of the Kyoto University Garden (KUG) on the first day of the experiment. The drawing beneath the photo is the site plan provided by Tiankai Liang. **C**. A participant is seated at the view spot in Murin-an. **D**. A participant seated at the viewing spot for the Kyoto University Garden (KUG). **E**. The average heart rate for all 16 participants during the 7 minute viewing period in the KUG (pink symbols) and Murin-an (blue symbols). Least squares regression lines are shown for both sets of points. **F**. The same data was normalized to the heart rate found in the first minute of viewing the garden.

For this second type of garden, we chose one on the main campus of Kyoto University (Figure 1B). The garden is located in the courtyard behind the civil engineering department building. The precise year of the garden’s construction and its designers are unknown. The garden features a stream at its center, with pine trees, shrubs, rocks, and a bridge carefully arranged around it. The garden also has a Himalayan cedar tree located at the edge of the stream, which was planted in 1912 by Sakuro Tanabe, a former dean of the school. At the end of the stream, a gazebo with a bench offers visitors a place to admire the view.

## Materials and Methods

Murin-an is a tourist garden which is quite famous for providing a sense of calmness to its visitors. The garden is open to the public year-round, except for administrative and weather-related closures. This presented a challenge to us as it was almost impossible to schedule a large enough block of time to complete our experiments. To illustrate this, consider that between 2016 and 2018, the average annual visitor count to Murin-an was 62,178, with more than 3,000 visitors per month. Because of the autumn colors, November is particularly a popular month for viewing. Because of these logistical difficulties in performing our experiments we had to negotiate a mutually acceptable time with the Kyoto city government and coordinate with the garden’s caretakers and staff. In the end, we had to perform our work in January on the one day that Murin-an was closed for its annual maintenance. While we had unfettered access to the facility, the lack of heat in the viewing room was an environmental factor over which we had no control.

### Spatial assessment

The space of the two gardens was surveyed using a VZ-400 3D laser scanner produced by RIEGL, plus a Canon E0S 600D camera with a Sigma 4.5 mm F2.8 EX DC Circular Fisheye lens. The VZ-400 uses a pulsed laser beam to scan the environment and generate a detailed 3D model. To ensure accuracy, the laser beam was positioned at the observer’s eye level (angle pitch 0.1°), and a 3D point cloud data was obtained with 5mm precision.

### Participants

The total number of participants in this experiment was 16, which was the maximum number we could use given the need to conduct our experiments in a single day at Murin-an. Among the 16 participants, 7 undergraduate students from Kyoto University Department of Civil Engineering – 5 males and 2 females – and 9 undergraduate students from Kyoto Art University Department of Environmental Design – 1 male and 8 females – took part in the study. All Kyoto University students were majoring in city planning and all of the art students were majoring in Japanese garden design. All participants were physically healthy with good eyesight (uncorrected vision or soft contact lens correcting no more than 0.7 in the decimal system, approximately 20/32).

Human research protocols for the study were approved by the Ethics Committee of Nagasaki University and all participants provided informed consent before data collection.

The study participants had two different educational interests. The nine art students from the Kyoto Art University studied Japanese gardens as part of their curriculum. The seven Kyoto University students were engaged primarily in policy and city planning courses and had limited detailed knowledge of Japanese gardens. Because it was on their campus, all policy/city planning students were familiar with KUG while the art students had mostly never seen it before. All policy/planning students visited Murin-an first and all art students visited KUG first. Thus, as part of the study design, all participants first visited a garden that they had never seen before.

### Eye-movement assessment

To record eye tracking and fixation, we utilized the EMR-10 eye-mark recorder (NAC Image Technology Co., Ltd. Japan), a device that captures high-speed video of eye movements using infrared light. It is non-invasive and uses a baseball cap with a small camera attached to the participant’s head via a headband. The camera records the eye positions and pupil movements. As the EMR-10 is designed to allow for head movement, participants can look around the garden with no restriction. To track the eye positions and pupil movements over time, the recorded data were analyzed using the company’s software, which aggregates gaze images within a specified time, converts them into a heat map, and automatically produces Excel files containing fixation information. We then analyzed eye-movement data using SPSS (version 19, International Business Machines Corporation), comparing mean values via repeated measure analysis of variance (ANOVA) using Bonferroni post hoc tests with one within-participant factor.

Significance was established at p < 0.05 (*) and p < 0.01 (**). Data are presented as means ± SEM.

### Heart rate assessment

We recorded participants’ heart rates to assess variability before and during the observation using a portable plethysmograph monitor (Iworx IWX/404 and PT 100 model). This equipment detects slight volume changes caused by blood pulsing through the finger vessels. Pulse rate data were continuously recorded throughout the experiment and analyzed using the software provided by the company, with the average pulse rate calculated every minute.

### Mood assessment POMS 2

The POMS 2 Brief form (DiLorenzo et al., 1999; McNair and Lorr, 1964; McNair et al., 1981; Nyenhuis et al., 1999) was used to measure mood changes during the experiments. The form is a self-administered adjective rating scale composed of 35 items with a 0–4 score and measures 7 mood states: anger, confusion, depression, fatigue, tension, vigor, and friendliness. The participants completed the POMS 2 Brief Form before and after observing the two gardens. As the participants’ native language was Japanese we used the Japanese translation of the questions (Heuchert et al., 2015)

### Questionnaire

In addition to POMS 2 Brief Form test, the participants were asked to answer the following questions: 1) Do you like this garden? 2) Have you been to this garden before? 3) Do you feel relaxed after viewing this garden? 4) Do you want to come again? 5) Have you visited any Japanese gardens before? 6) If yes, how often do you visit a Japanese garden? 7) Any other comments?

### Garden observation

The experiment at Murin-an was scheduled for January 18, 2023, while the one at Kyoto University Garden (KUG) was scheduled for January 17 and 19, before and after the Murin-an experiment. Half of the participants were scheduled to observe KUG on January 18 and the other half on January 19. As discussed above the logistics of access to the Murin-an garden meant that we were restricted to a single day in January. The temperature on all three days was 6°C at 9:00 a.m., 12°C at 12:00 noon, and 13°C at 3:00 p.m., with partially cloudy weather and low wind speeds. Participants were provided with a warmer and blanket during their viewing of the KUG.

Participants arrived at a waiting station where they filled out a form. They were then escorted to the garden station (Figure 1C, D), which was equipped with a viewing seat, table, and movable partition to block the view while attaching and calibrating monitors. After arriving at the garden station, participants were instructed to perform the following tasks:

1. Fill out the POMS 2 form while sitting with their backs to the garden.
2. Wait while the electrocardiograph and eye tracking monitors were connected by the research assistant.
3. Rest for 3 minutes with no view of the garden.
4. Turn around and observe the garden for 7 minutes.
5. Turn back around and sit with their backs to the garden.
6. Complete the post-exposure POMS 2 form and the questionnaire.
7. Wait while the electrocardiograph and eye tracking monitors were removed by the research assistant.

## Results

### Spatial measurements

The dimensions of the two gardens were similar. The Murin-an garden measured approximately

85.5 m in depth and 36.6m in the perpendicular direction while the Kyoto University courtyard where the garden was located measured approximately 84.5 m in depth and 32.0m in the perpendicular direction. The horizontal and elevation angles from the viewing point were also similar, with Murin-an having a horizontal angle of 23.68° and elevation angle of 9.9°, and KUG having a horizontal angle of 45.29° and elevation angle of 8.2°. The high visibility horizontal viewing angle is 50.97° and the maximum elevation angle is 17.2°.

### Heart rate changes

We have previously measured the response of participants’ heart rate when viewing a Japanese style garden (Goto et al., 2017; Goto et al., 2014; Goto et al., 2013; Goto et al., 2018) and the responses of these participants were consistent with our earlier findings. Before the garden observation, participants had a 3-minute resting time during which their average pulse rates were 74.2 bpm in KUG and 79.5 bpm in Murin-an. Though this difference is notable as the participants were the same for both gardens, the reason for it is unknown. During the 7-minute viewing period the average heart rate in KUG increased by 5%, while that in Murin-an decreased steadily (Figure 1E). The difference in the two trends is clear and a linear regression fit to the two sets of data points have significantly different slopes. As the baseline heart rates varied in the two gardens, we replotted the data as the change from the heart rate observed during the first minute of the session (Figure 1F). Calculated in this way, the differences in between the two gardens were significant at every 1-minute bin.

### Eye movement

The heart rate data shows that, as in our previous studies, viewing a well-constructed garden can impact the physiology of the viewer within minutes. We next sought to investigate the eye movements of the participant for clues as to what features of the visual field were predominant in eliciting this effect. Figures 2A and 2B show a heat map of the eye fixation points of all 16 participants on the 1200 x 675 pixel picture pane. This summary map makes several clear points. The first is that the participants primarily focused on the center of the visual environment in both gardens. Second, the fixation points were far more distributed in Murin-an than in KUG. The control KUG garden had many of the same visual elements found in Murin-an – water, stones, trees, a bridge – but we were concerned that participants would be distracted by the geometric features of the building to their right. This worry proved unnecessary as very few of the fixation points localized to the building (Figure 2B). In Murin-an by contrast, despite the absence of any obvious feature to attract the participants’ gaze, their fixation points distributed over the entire field of view including the edge (Figure 2A). In addition to the difference in their distribution, quantification of the data revealed additional significant differences during the 7 minutes of observation. When observing the Murin-an garden, the total number of fixation points (6,620) was greater than when viewing KUG (5,885 – *p=*0.01). But during any given minute, while the average number of fixation points was elevated in Murin-an compared to the KUG (Figure 2C), the difference between the two gardens only trended higher (Figure 2D – *p* = 0.069 by unpaired t-test). Far more dramatic were the differences between the two gardens in the distance covered by the shifting gaze of the participants’ (Figure 2E), and speed of movement between fixation points (Figure 2G). This difference was true from the very beginning, did not change during the 7 minutes viewing period, and the difference between the two curves was highly significant (Figure 2F, H – *p* < 0.001 for both graphs).

**Figure 2.**
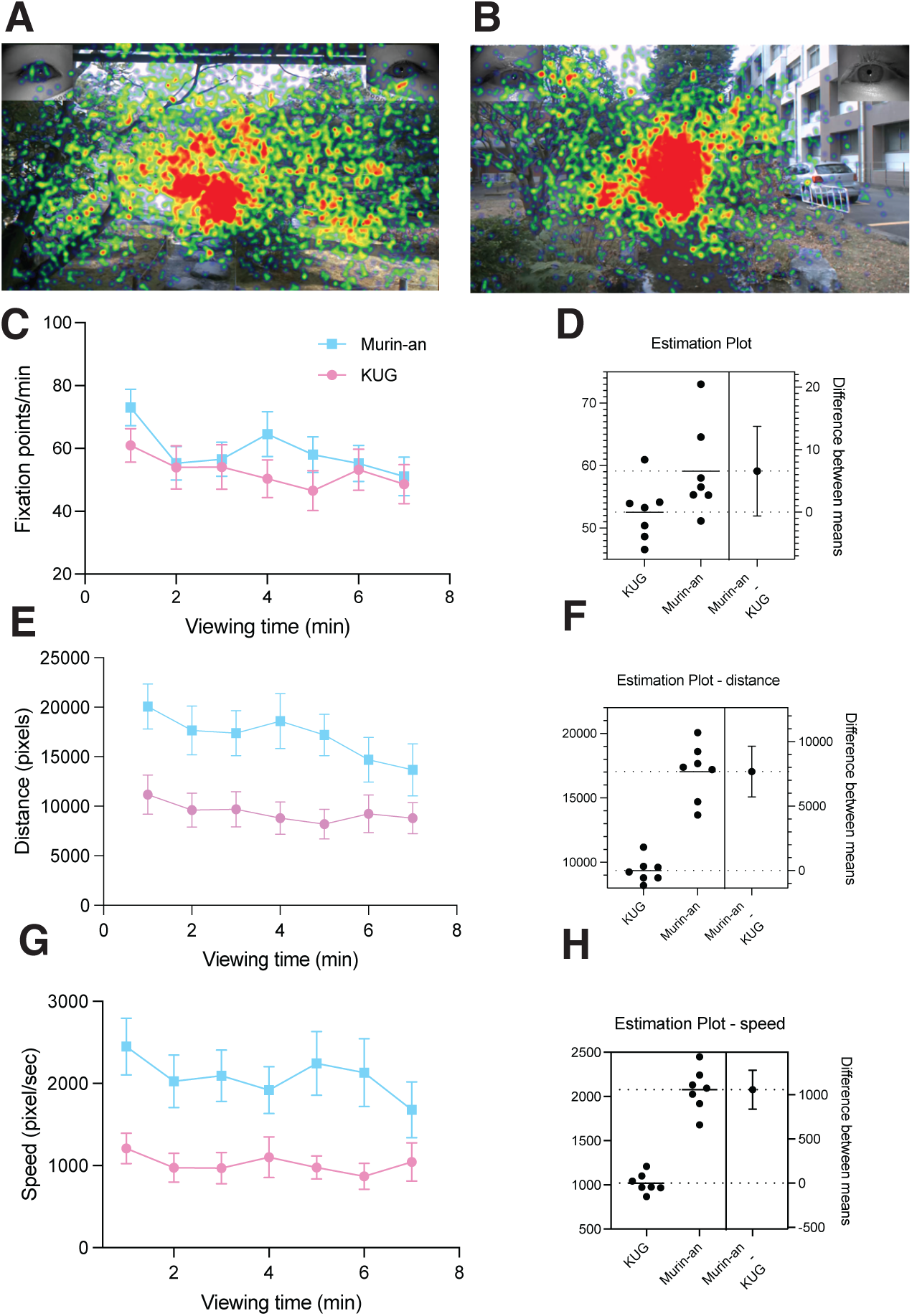
Heat maps of the fixation points of all 16 participants during their 7-minute viewing of the Murin-an (**A**) and KUG (**B**)gardens. The color scale is graded from blue (fewer fixation points) to red (more fixation points). **C**. The average number of fixation points during each one-minute time bin of the view period in the KUG (pink symbols) and Murin-an (blue symbols). **D**. Estimation statistics showing the effect size and the distribution of the differences. **E**. The average distance traveled by gaze of the 16 participants during each minute of the 7-minute viewing session was greater in Murin-An than in the KUG (*p* < 0.0001). **F**. Estimation statistics showing the effect size and the distribution of the differences. **G**. The average speed of the participant’s eye movements during viewing session was greater in Murin-an than in the KUG (*p* < 0.0001). **F**. Estimation statistics showing the effect size and the distribution of the differences.

To further analyze the relationship between the fixation points and observed objects in the two gardens, we divided the picture pane into 15 blocks based on their content (Figure 3A, B, E). We calculated the number of fixation points in each block and normalized this value to the fraction of the entire visual field represented by that block. In KUG (Figure 3B, D), the object most focused on was the bridge, followed by middle ground plants, and water. In Murin-an (Figure 3A, C), the most noticed blocks were the water, followed by what we termed the middle ground and the bridge. Note that these high-interest objects are arranged differently in the two gardens. In the KUG the three most-focused objects were all in the center of the visual field while in the Murin-an garden, the objects were in a more horizontal arrangement. Note also that the size or type of object did not determine its focus; instead, it was the object’s position within the overall spatial composition that guided the gaze. Most likely, in KUG, the bridge’s location at the vanishing point of the picture pane directed the gaze toward it while in the Murin-an garden, we believe that the river’s acute angle in the center directed the gaze from side to side, eventually landing on the bridge.

**Figure 3.**
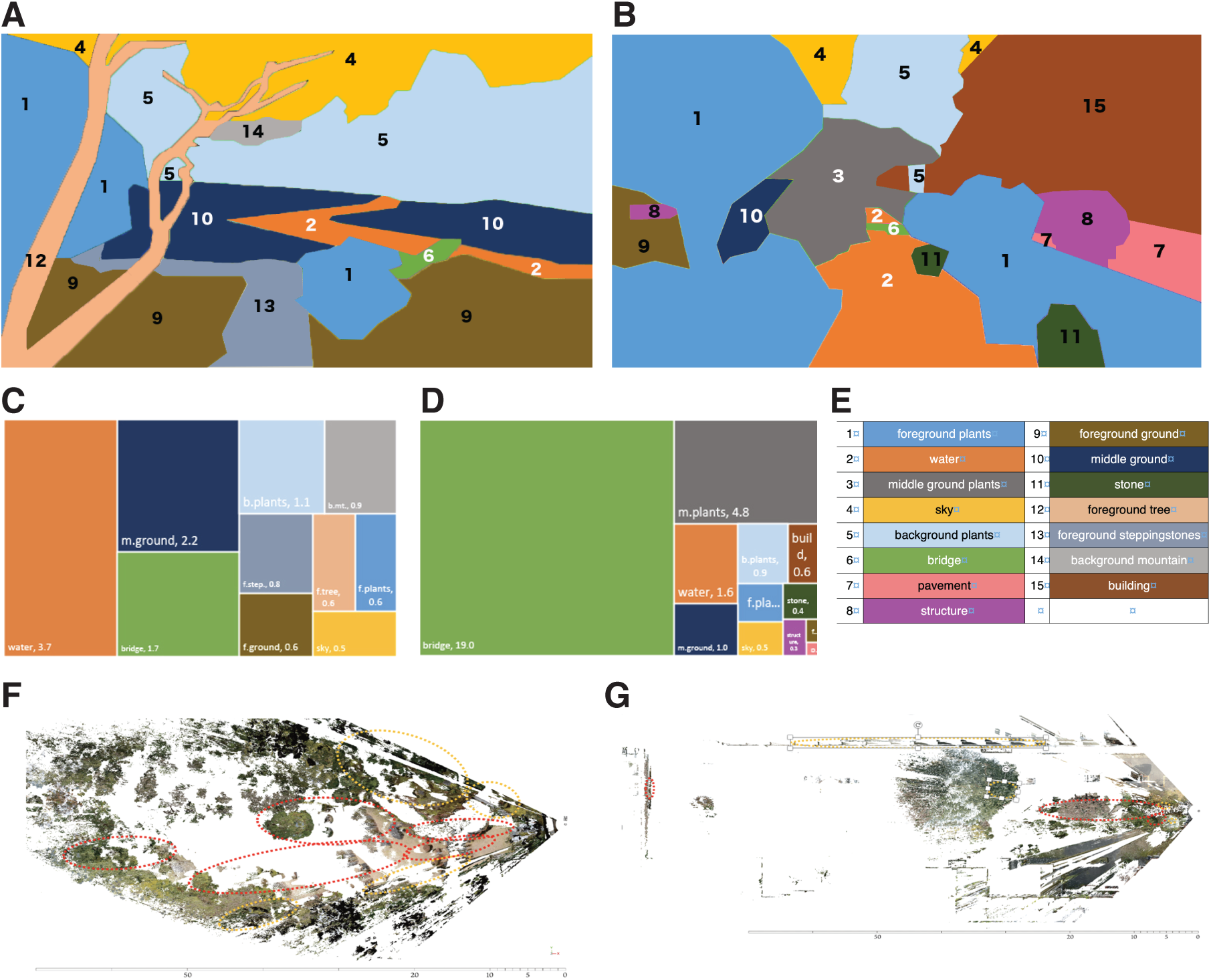
The assignment of content blocks to the visual scene in the Murin-an garden (**A**) and the KUG (**B**). The times that the focal point of all 16 viewers were within the content blocks shown for the Murin-an garden (**C**) and the KUG (**D**). For each block the percentage of time spend in each region was normalized to the percentage of the total visual field area covered by that block. **E**. The color code for panels A-D. We calculated the distance of each focal point from the viewer (the Z-dimension) and plotted these points as shown for Murin-an (**F**) and for the KUG (**G**). The plot reveals the focal points in the two gardens as if viewed from above.

Our analysis enabled us to determine not only the 2-dimensional (X-Y) coordinates of the viewers gaze, but also its depth (the X-Z coordinates – Figures 3F and G). This figure emphasizes the dramatic differences in the ways in which the participants viewed the two gardens. In the KUG, not only did the viewer tend to focus on the center of the field in the X-Y plane, but they were also focused primarily on the nearer objects in the X-Z plane. The eye-movement fixation points in Murin-an were distributed throughout the background, middle ground, and foreground. In the KUG by contrast, most fixation points were on the plants in foreground, with only 2% on the background plants. In 3-dimensional space therefore, the participants in the KUG attended mostly to the lower areas and to the front and middle depths, while in Murin-an the attention was on the upper areas and far more broadly distributed through the entire depth of the visual field. These differences may have been due in part to the large Himalayan cedar tree in the KUG blocked the view of distant scenery. In the Murin-an garden, as intended by its designer, there is an open view of Mt. Higashi, located several kilometers away.

### Qualitative assessments

#### Questionnaire and POMS

The two gardens elicited very different qualitative responses in the participants. Participants’ responses to a simple questionnaire (see Methods) showed that they liked (*p* = 0.0001), felt relaxed in (*p* = 0.007), and wanted to revisit (*p* = 0.001) the Murin-an garden significantly more than the KUG (Figure 4A). The POMS test results in the two gardens also differed. The POMS evaluates seven mood states—anger (AH), confusion (CB), depression (DO), fatigue (FI), tension (TA), vigor (VA), and friendliness (F). The last two states, vigor and friendliness, did not have obvious relevance to our study but they are integrated into the questions and could not be removed without validation. To determine whether viewing the garden impacted mood, we administered the POMS both before and after participants viewed the two gardens. The scores of the first test administered (before the garden was viewed) varied considerably among the participants. To represent the data therefore, we calculated the responses of each individual participant as the difference between their responses on the second test (after viewing) and the first (before viewing). We then compared the magnitude of this after-minus-before difference in the KUG with the magnitude of the difference in the Murin-an garden (Figure 4B). In the POMS, a positive value indicates an improvement in mood. As represented in Figure 4B, however, if the effect was larger in the Murin-an garden then the value reported for the difference of the difference will be negative (i.e., if Murin-an > KUG then KUG - Murin-an will be less than zero). This is precisely what we saw; the Murin-an garden had a stronger effect on the POMS score than the KUG. For the first five measures of mood, the difference in Murin-an was greater (reflected in a negative value in the graph). For measures of the vigor and friendliness components of mood, there was virtually no change in either garden (not shown) and thus the inter-garden comparisons were insignificantly changed. Overall, a two-way ANOVA of the results suggests a highly significant difference between the two gardens (p < 0.001).

**Figure 4.**
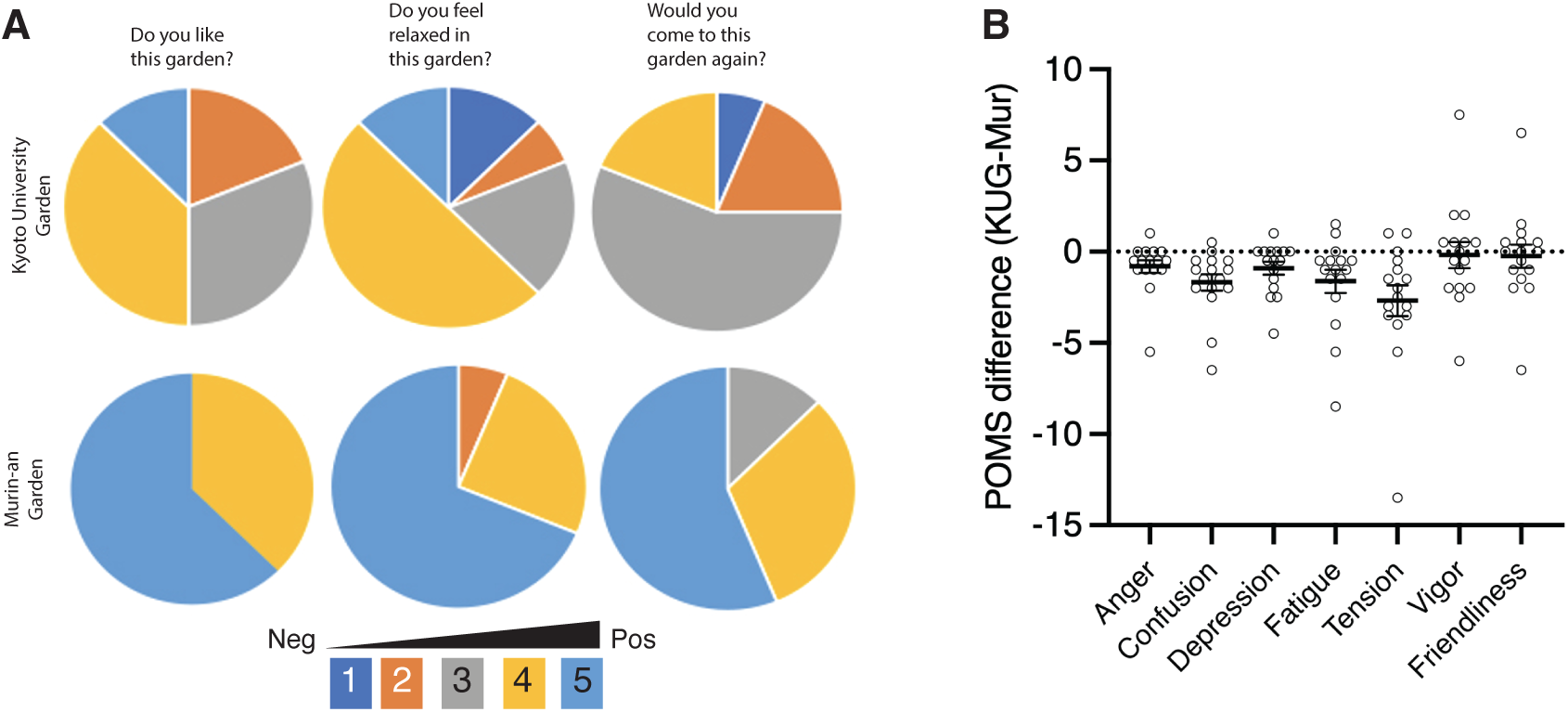
**A**. The response of the viewers to three of the questions asked on the short questionnaire. The size of each pie wedge is proportional to the percentage of respondents who assigned that score to their answer. The legend at the bottom shows the values of the 5-point scale used. **B**. The results of the POMS instrument to detect the mood of the subjects before and after garden viewing. The X-axis represents the seven categories of mood, the Y-axis represents the difference of the differences. For each subject, we measured their responses on the POMS before and after the 7-minute garden viewing session. A positive value of the after-minus-before difference indicates an improved score on that aspect of mood. We repeated this measure for each subject in each of the two gardens. To compare the effects of two gardens on mood, for each subject we subtracted their after-minus-before difference in the Murin-an garden from the after-minus-before difference in KUG. The mostly negative scores of this metric indicate that the improvement of mood in Murin-an was greater than in the KUG.

### Impact of student background on the garden response

The backgrounds of the 16 participants differed in ways that led us to ask whether these differences affected their responses to the garden in any consistent way. The seven students from Kyoto University were from a program in policy and city planning. While they were familiar with the importance of green spaces, of the seven, two had never visited a Japanese garden, four visited a few times a year, and only one visited a few times a month. By contrast, all nine participants from Kyoto Art University were studying to become Japanese garden specialists and had taken classes on Japanese gardens at the university. Seven reported that they visited Japanese gardens several times a month; two reported visiting a few times a week. Further, as it was on their campus, all seven policy/city planning students had seen the KUG previously, but only two had been to Murin-an before the experiment. By contrast, of the 9 art students, only one had visited the KUG, but four had previously visited Murin-an.

Each of the four groups showed a unique heart rate response to the gardens. Once they had settled down during the pre-viewing period in the KUG, the heartrates of the students with a background in Japanese garden techniques climbed steadily throughout the garden viewing session (Figure 4C, dark red). This was suggestive of an increased level of stress/irritation. In the Murin-an garden, these same students had an initial rise in their heartrate during the first minute of viewing followed by a slow but steady decline in rate (Figure 4C, dark blue). The policy/city planning students, with less previous exposure to Japanese gardens, responded quite differently. In the University setting, they were quicker to settle down during the pre-viewing period and once they were allowed to view the KUG, their heart rates remained virtually unchanged during the viewing period (Figure 4C, light red). In the Murin-an garden, their resting rate was noticeably higher than either their own rate before viewing the KUG or the rate of the art students in Murin-an. Despite this suggestion of elevated stress/anxiety, once the viewing period began, their heartrates also decreased and remained low throughout (Figure 4C, light blue).

Overall, we noticed that the absolute heartrate could vary but the relative change over the viewing period was consistent in the two different groups – declining more in the Murin-an garden than in the KUG. The slight increase that we noted earlier in the KUG heartrate (Figure 1E, F) appears to have been driven almost entirely by the Kyoto University Art students (Figure 5A, dark red). We also asked whether the viewing strategies of the two groups – total eye movement distance and speed of movement –differed in the two different gardens. From minute to minute there were more noticeable fluctuations in the distance traveled (Figure 5B) and speed of the eye movements (Figure 5C) in the Murin-an garden in both groups of students, compared with the KUG where both measures of gaze remained relatively stable and constant during the viewing period.

**Figure 5.**
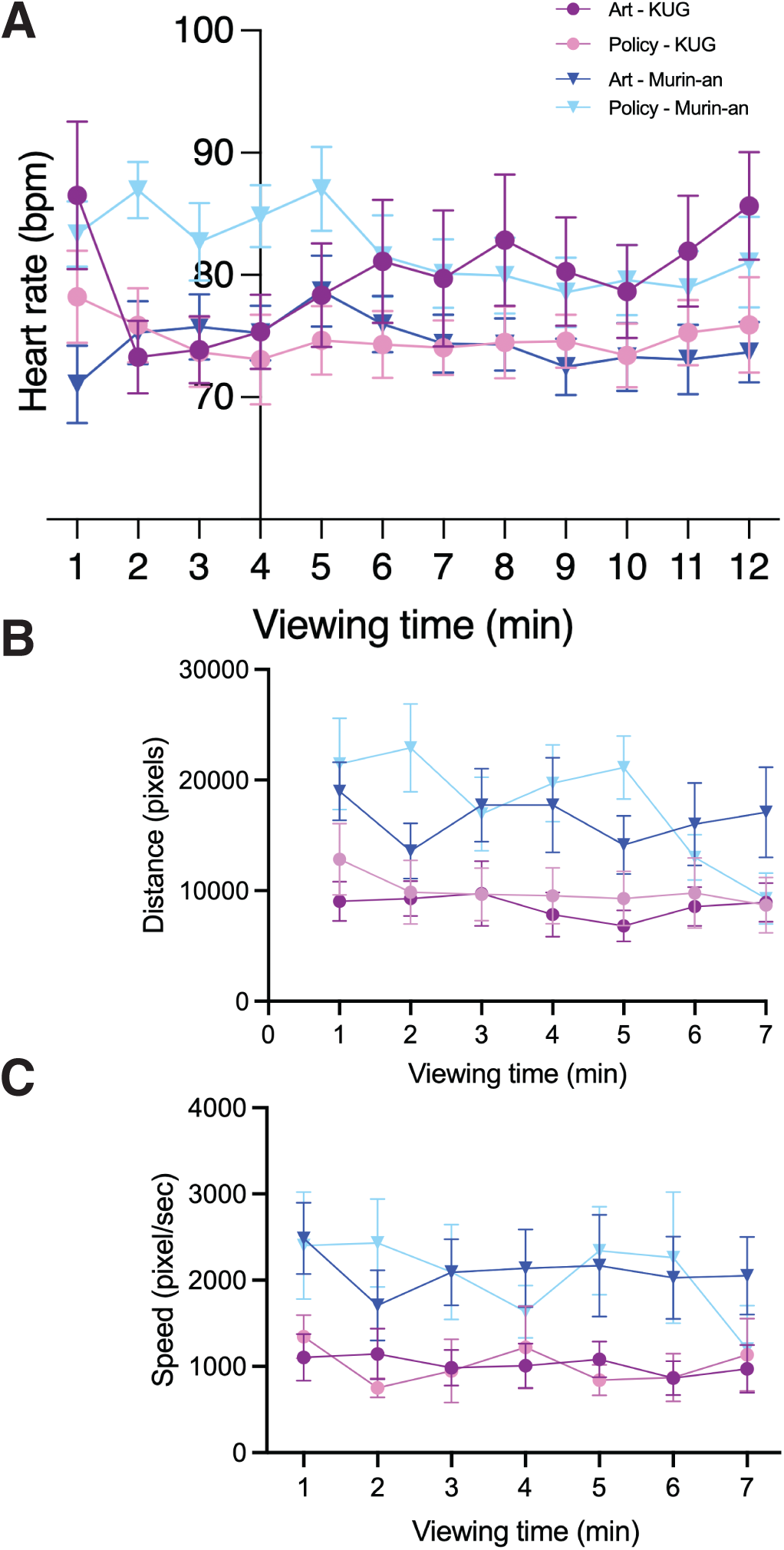
**A**. The heart rate of the participants in the two gardens (Murin-an and KUG) plotted separately for the two groups of students (policy/planning and art) for the entire session including the 3-minute pre-viewing period. **B**. The average distance traveled by the gaze of the 16 participants during each minute of the 7-minute viewing session plotted separately for the two groups of students. The identity of the lines is the same as indicated by the legend in panel A. **C.** The average speed of the shifting gaze of the 16 participants during each minute of the 7-minute viewing session plotted separately for the two groups of students. The identity of the lines is the same as indicated by the legend in panel A.

The qualitative observations also differed between the two groups, but not dramatically. In Murin-an scores for the question about feeling relaxed by viewing the garden increased by 124% among the city planning students and 140%, among the art students compared to KUG. Similarly, the scores for the question about if they want to revisit the garden increased by 122% and 153% among planning and art groups, respectively in Murin-an compared to KUG (Figure 5). Overall, Murin-an was more popular than KUG among all participants, with art students evincing a stronger preference for Murin-an than the city planning students. We performed a 2- way ANOVA (two gardens, pre-post) for the POMS results of the two groups. For the art students, we found a significant difference between the pre and post values for both gardens (F(1, 8) =6.98, p<.05) as well as a trend between the pre-viewing values of the two gardens. For the policy/planning student we found a significant difference between pre and post values in Murin- an, but no change in the KUG. In short, the art students’ mood improved in both gardens, while policy students’ mood improved significantly only in Murin-an.

## Discussion

Our findings provide several important advances in our ongoing studies of Japanese gardens and their effect on the viewer. Our early work in this area focused on elderly participants, including those with advanced dementia (Goto and Fritsch, 2011; Goto et al., 2014; Goto et al., 2013). These studies revealed that there were features in the elements and design of a Japanese style garden that were unique in their ability to engage participants and lower both their qualitative sense of tension and its physiological manifestation in heart rate. Perhaps most intriguing was how viewing the garden cut through the fog of dementia and produced effects, even in participants whose MMSE (Mini-mental status exam) scores were below 15 (Goto et al., 2018). The robust nature of the effect led us to explore its neurobiological underpinnings, which led us to an interest in tracking eye movements (Goto et al., 2017; Goto et al., 2020; Liu et al., 2020). Our hypothesis was that, by correlating the object of a participant’s attention with the momentary change in their heart rate, we could identify those features of the garden – a stone lantern, a water element, a specific type of plant – that were most responsible for eliciting the calming effect we had documented. The results did not support this idea but did reveal that the pattern of eye movements differed significantly when participants viewed a Japanese style garden (Goto et al., 2020).

The current study offers a possible explanation as to why our hypothesis was incorrect, and ironically, it may have been the uncomfortable nature of the viewing environment that was the key to this insight. We chose to measure the effect of the Murin-an garden because it is a classic and highly regarded design that we hoped would maximize the signal we were seeking (a more detailed description of its history and design principles can be found in the Supplementary Materials). Unfortunately, it is also a popular tourist attraction and because of this, our access for the purpose of experimentation was restricted to a single day in January. Kyoto winters are cold and the villa from which the garden is viewed is not heated. The KUG viewing spot was completely outdoors and the viewing sessions were held during the same January days. We provided blankets for warmth at both locations, but most participants remarked about the conditions. The wintertime viewing schedule also meant that both gardens were devoid of any flowering plants and their deciduous trees were bare. The subdued colors meant that the participants were influenced primarily by the design features of the two gardens.

The pulse rate measurements show the effects of this minimalist stimulation. In previous work, in comfortable conditions, even a less than professional Japanese style garden was sufficient to induce a perceived and physiologically detectable calming effect (Goto et al., 2013). Yet viewing the KUG led to no significant drop in heart rate, even though the POMS revealed that most measures of mood improved. Despite the cold, however, the Murin-an garden was highly effective and produced a calming effect on both heart rate and mood. Both groups of students experienced an increase in their heart rate during the first minute in Murin-an after which the policy/city planning students showed a steeper decline in heart rate while the art students heart rate decreased more slowly. In other words, viewing Murin-an had a greater calming effect on the participants who had less knowledge of Japanese gardens. This suggests that the relaxation effect of Murin-an is much higher than that of a typical Japanese garden and also more effective for first-time viewers.

The eye-tracking data add an important dimension to the analysis. We found a substantial difference in the gaze movements during the viewing sessions of the two gardens. We found that to some extent, a participant viewing a garden they had never seen before led them to take in a wider range of views. While this may have been a factor for some participants, this effect was minor compared to our observation that the range of fixation points in all three planes, the distance traveled by the eyes, and the speed with which they moved were all significantly greater in Murin-an than in the KUG. As we observed previously (Goto et al., 2020), the participants’ gaze in both gardens tended to concentrate in the center of the visual field. Nonetheless, the distribution of points was far broader in Murin-an, extending to the edge of the visual field especially in the horizontal direction. In the KUG, even the regular geometry of the windows in the building at the right edge of the visual field did not draw the participants’ attention.

We note the analogies between our findings and the principles used in a technique known as eye-movement desensitization and reprocessing (EMDR). EMDR was developed by psychologist Francine Shapiro who is credited with first noting the association between eye movement and mood (Shapiro, 1989; Shapiro et al., 1994). The theory of the method is to reduce stress by providing bilateral stimulation through the manipulation of eye movement during the recollection of a specific memory. After its initial discovery other sensory domains such as sound and touch have been added (Shapiro, 1999). The technique is widely used, especially to treat the persistent stress associated with traumatic events (e.g., PTSD), but its neurological basis is not understood (Pierce et al., 2023; Valiente-Gomez et al., 2017; Zarghi et al., 2013). The success of the technique has been suggested to be due in part to the strong bilateral nature of the visual stimuli used and some authors have noted its relationship to the REM phase of sleep (Stickgold, 2002).

Viewing a garden from a single vantage point is in no way equivalent to a session of EMDR and yet the work in this area and our own observations align in ways that offer suggestions for future exploration. For EMDR, the finding that the effect of the Murin-an garden was associated with increased focal points in all three dimensions suggests that when visual stimuli are used it may prove worthwhile to alter the distance of the stimulus as well as its left-right position. This suggestion receives additional support from our previous finding that viewing a flat projected image of a garden is less effective than exposure to the full 3-dimensional scene (Sun et al., 2018). For our own studies, the strong effect of rapid bilateral visual stimulation that is achieved during EMDR suggests that it is the wider, more rapid, eye movements that are elicited in a well designed garden such as Murin-an that are the reason for the physiological and psychological effects that we observe. Thus, our search for a specific item in the garden that acts as the source of relaxation was always going to fail. Instead, it suggests it is the totality of the design, and the rapid and broad eye movements evoked, that are the key ingredients. The later EMDR work of Shapiro and others showing the effectiveness of other sensory modalities urges more consideration of the sonic and even olfactory environment of a garden in eliciting the full stress reduction effects that we observe.

## Supporting information

Goto et al - Supplemental Material

## Notes

### Competing Interest Statement

The authors have declared no competing interest.

