## Supplementary material for "Correlating stress reduction and eye movement patterns in a world famous Kyoto Japanese garden": Goto et al - Supplemental Material

### Supplemental information

#### An architectural and historical account of the two gardens used in this study.

Both Murin-an and KUG are ornamental Japanese gardens centered on a body of water meant to be viewed from a single vantage point that provides a controlled perspective of the scenery. Their design concepts and management, however, differ considerably. Murin-an was designed to recreate the natural scenery that its owner, Yamagata Aritomo, would have seen in Hagi, his birthplace. It is a picturesque garden that draws inspiration from Hagi's natural beauty, particularly its mountains and rivers (Figure S1A). Yamagata designed the garden with a river flowing through a field a view of Mt. Higashi in the background. Although the garden is small, Yamagata wanted it to be viewed as an extension of Mt. Higashi, creating the impression that the river water was flowing from there. In contrast to the traditional Japanese garden style of placing stones vertically, Murin-an features large stones laid horizontally to avoid obstructing the line of sight to the mountain from the sitting point. The river is designed in a zig-zag pattern that naturally directs the viewer's gaze diagonally across the garden, creating a sense of expansion for viewers. A small waterfall is also included to produce the sound of water near the sitting point, ensuring that this sound reaches the viewer's ears.

Despite being currently surrounded by buildings and densely populated (Figure S1B), the Murin-an garden has preserved Yamagata's original design concept through constant management and attention to detail. For example, the height and density of trees in the background are regulated to conceal the buildings behind and create a sense of depth, extending the view of Mt. Higashi. Although the Murin-an garden is triangular with two sides bordering the road, the view of adjacent buildings and roads is completely obscured by the trees in the garden. In the foreground, the lawn is adorned with wildflowers and various types of moss, meticulously maintained by hand to create a velvety green carpet without any gaps. The water flowing across the garden's field glows in the middle ground, and the sound of water falling on the steps immediately next to the viewer may be realistic enough to be mistaken for the distant river. The garden is immaculately clean, and its well-maintained details add complexity and depth to the space. As one spends more time at Murin-an, its captivating and awe-inspiring qualities become increasingly apparent. Murin-an seems to exemplify Kaplan's theory of the effect of unconscious mild aesthetic experiences in nature (ART).

By contrast, KUG is enclosed by school buildings with a commemorative cedar tree, planted years ago by the second dean of the School of Engineering, serving as its focal point (Figure 1B). The cedar has grown into a large tree and the school buildings and asphalt roads are visible on both sides of the garden, with cars parked on the road. Unlike Murin-an, KUG has no management scheme to ensure the maintenance of the overall balance in the garden's original design. When viewing the garden, one's gaze is drawn to the large canopy of the cedar, which dominates the field of vision. The course of the river and buildings on the side guide the gaze straight to the center of the field of vision – the center of the river. This combination of distractions makes it difficult to associate the water and its few stones with a natural flowing river. Instead, the more one looks at KUG, the more one notices stagnant water and discarded items that do not belong.

Japanese gardens are more than just the planting of pruned pine trees and setting small stones; they evoke an emotional response and establish a personal connection with nature. The effectiveness of a Japanese garden depends not only on its design concept but also on its maintenance. This is crucial for conveying the garden's message and can make or break its impact. In Murin-an, the eyes move faster, stopping at more points in a wider range of vision and depth, inspiring a connection with the beautiful natural surroundings. By contrast, looking around KUG, one is constantly reminded of negative elements, such as stagnant water and detritus, instead of being reminded of and inspired by the beauty of nature.

In this experiment, we found that Murin-an garden had a calming effect on all participants, both psychologically and physiologically, within 3 minutes. This is the effect that attracts and heals countless global visitors, rendering this garden famous. Many famous Japanese gardens built in temples typically reflect the ideas of Buddhism; however, the concept behind Murin-an is unusual in that it was designed to showcase scenery that is familiar to everyone. Whether Murin-an is particularly relaxing because the landscape symbolized there is one that everyone can relate to is not known. These thoughts motivate us to explore whether the effects of viewing Japanese gardens differ depending on their design concept.

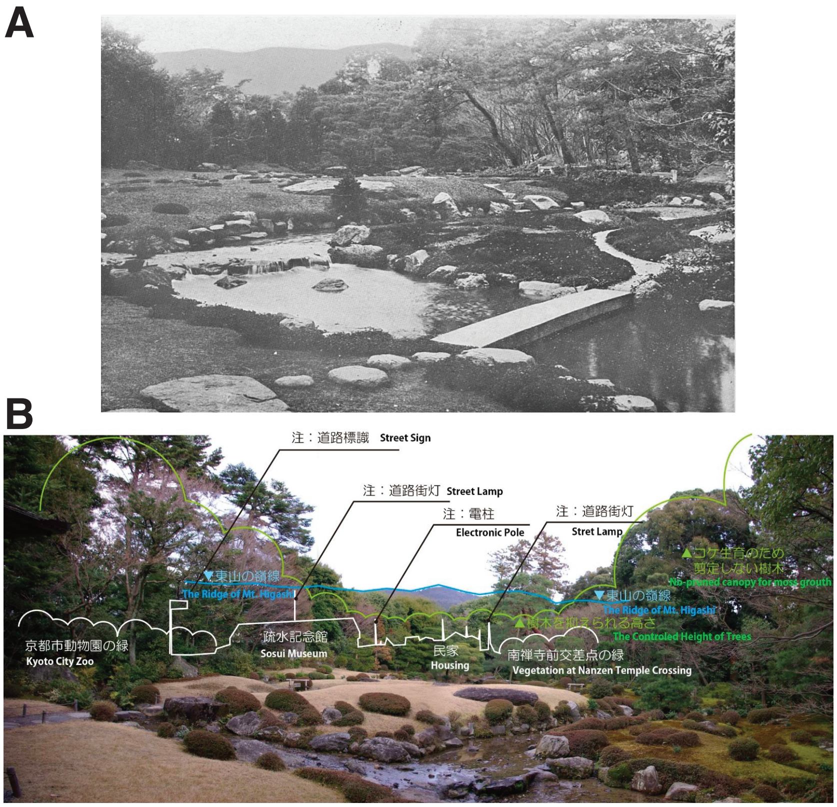


Figure S1.
**A**. A photograph of the Murin-an garden in its original form (from Tamura, 1935). **B**. A recent photograph of Murin-an annotated with the features of the environment that are hidden when viewed from the specified viewing point in the villa (white lines and labels) and the adjustments that have been made to retain the original spirit of the design. The blue lines illustrate the ridge containing the distant Mt. Higashi.
